# Effects of Gallic Acid on Antioxidant Defense System and Nrf2 Signaling in Mice with Benzene-Induced Toxicity: In Vivo, In Vitro, and Computational Study

**DOI:** 10.1101/2025.09.04.674095

**Authors:** Toba Isaac Olatoye, Blessing Augustine Afolabi, Michael Odunayo Ogundipe

## Abstract

**Background:** Benzene exposure is a well-known cause of toxicity in several tissues, primarily through the generation of reactive oxygen species (ROS) and disruption of redox homeostasis in the bone marrow. The resulting oxidative stress impairs hematopoiesis, weakens antioxidant defenses, and increases cellular damage. The transcription factor Nrf2 plays a central role in counteracting oxidative stress by regulating the synthesis of antioxidant and detoxifying enzymes, however, its activity is tightly controlled through Keap1-mediated degradation. Antioxidant therapy, particularly with the use of phytochemicals like gallic acid, has emerged as a promising strategy to mitigate these effects. Although the antioxidant potential of gallic acid is well documented, there is still limited integrative evidence regarding its molecular mechanism in counteracting oxidative stress.

**Objective:** This study aims to examine the protective effects of gallic acid on antioxidant defenses, oxidative stress biomarkers, and hematological parameters in mice with benzene**-**induced toxicity, while also evaluating its potential to modulate Nrf2 signaling through molecular docking.

**Methods:** Thirty-six mice were randomized into six groups of six animals each. Group A served as the normal control, while the remaining groups (B-F) were orally administered benzene (150 mg/kg body weight) for fourteen days. Groups D, E, and F received simultaneous oral administration of gallic acid at 25, 50, and 100 mg/kg body weight, respectively. Group C received ascorbic acid (50 mg/kg body weight) as a reference, while Group B was not treated and served as the negative control. Biochemical analyses of tissues (erythrocyte, heart, liver, kidney, femur, and spleen) were performed to assess antioxidant enzyme activities (SOD, CAT, GPx, GST), the non-enzymatic antioxidant GSH, and levels of oxidative stress biomarkers (MDA, PCO, NO, PC). Toxicity in hematological parameters were determined from whole blood, while molecular docking was used to evaluate the binding affinity and interactions of gallic acid with the Kelch domain of Keap1.

**Results:** Exposure to benzene significantly reduced antioxidant enzyme activities, depleted GSH and PC, increased MDA, PCO, and NO levels, and altered hematological parameters (WBC, RBC, HGB, HCT, PLT, LYM) in untreated mice at *P<.05*, which was consistent with oxidative and nitrosative stress. In contrast, treatment with gallic acid significantly restored antioxidant enzyme activities, increased GSH and PC levels, reduced the concentrations of MDA, PCO, and NO, and improved hematological parameters in a manner comparable to ascorbic acid at *P<.05*. Molecular docking also revealed strong binding affinity (binding energy: -6.8 kcal/mol) and promising interactions of gallic acid within the Keap1 Kelch domain, suggesting a potential mechanism for Nrf2 stabilization and nuclear translocation.

**Conclusions:** Our findings demonstrate that gallic acid improves the cellular antioxidant defense system and provides protection against benzene-induced oxidative stress in mice. In addition, it may also activate Nrf2 signaling by disrupting the Keap1-Nrf2 complex, thus promoting cellular resilience against oxidative stress.

## Introduction

Benzene is a volatile aromatic hydrocarbon widely used in many industrial applications and is also recognized as a potent environmental and occupational toxicant. Chronic exposure to benzene, even at low concentrations, has been associated with myelotoxicity, hematopoietic toxicity, immunosuppression, and an increased risk of leukemia and other blood disorders [1–6].

A principal mechanism underlying benzene-induced toxicity is oxidative stress, which arises from the overproduction of reactive oxygen species (ROS) and reactive nitrogen species (RNS) during benzene metabolism, especially in the liver and bone marrow [7]. Oxidative stress has been implicated either as a primary cause of metabolic disorders such as atherosclerosis, paraquat poisoning, radiation-induced pneumonitis, and fibrosis, or as a secondary contributor to the progression of diseases including chronic obstructive pulmonary disease (COPD), cancer, diabetes, hypertension, cardiovascular and neurodegenerative diseases [8].

In eukaryotes, the antioxidant defense system plays a critical role in maintaining redox homeostasis by neutralizing excessive ROS and RNS, thereby preventing oxidative damage. This system comprises enzymatic antioxidants such as: (i) superoxide dismutase (SOD), which catalyzes the dismutation of reactive superoxide radicals (O_2_^•−^) into hydrogen peroxide (H_2_O_2_) and oxygen (O_2_); (ii) catalase (CAT), which decomposes H_2_O_2_ into water (H_2_O) and O_2_, thereby preventing hydroxyl radical (^•^OH) formation; (iii) glutathione peroxidase (GPx), which reduces H_2_O_2_ and lipid hydroperoxides into water and alcohols respectively, using reduced glutathione (GSH) as a cofactor; (iv) glutathione S-transferase (GST), which catalyzes the conjugation of GSH to electrophilic compounds like benzene oxide (a reactive intermediate formed during benzene metabolism by hepatic cytochrome P_450_ enzymes) for detoxification; and (v) glutathione reductase (GR), which regenerates GSH from its oxidized form (GSSG), in order to maintain the cellular pool of GSH. Disruptions in the activities of these enzymes often lead to an accumulation of oxidative stress biomarkers such as malondialdehyde (MDA), a marker of lipid peroxidation; protein carbonyls (PCO), which indicate oxidative damage to proteins; nitric oxide (NO), which contributes to nitrosative stress by forming peroxynitrous acid (ONOOH) (a potent and highly indiscriminate oxidant) upon reacting with superoxide radicals; and the depletion of the most abundant non-enzymatic antioxidant GSH and cellular protein concentration (PC) [8–11].

At the molecular level, the cellular response to oxidative stress is predominantly regulated by the transcription factor nuclear factor erythroid 2-related factor 2 (Nrf2). Under physiological conditions, Nrf2 is tightly regulated in the cytoplasm by Kelch-like ECH-associated protein 1 (Keap1), a substrate adaptor for the Cullin-3 (Cul-3) E3 ubiquitin ligase complex. Keap1 binds to Nrf2 through two motifs in the Nrf2-ECH homology 2 (Neh2) domain: a high-affinity ETGE (Glu-Thr-Gly-Glu) motif known as the “hinge” and a lower-affinity DLG (Asp-Leu-Gly) motif also referred to as the “latch”, thus, targeting Nrf2 for degradation by the ubiquitin proteasome system (UPS) [12]. This reaction ensures cellular Nrf2 levels remain low in the absence of oxidative stress. However, during oxidative stress, reactive cysteine residues in Keap1 undergo covalent modifications or conformational changes, making Nrf2 to dissociate from it and escape UPS degradation. Stabilized Nrf2 resulting from this process then translocate into the nucleus, where it forms heterodimers with other transcriptional regulatory proteins. These protein complexes bind to the antioxidant responsive element (ARE), and activate the transcription of genes responsible for antioxidant defense and xenobiotic detoxification [13–14].

Almost all known ARE activators act as indirect inhibitors of the Keap1–Nrf2 interaction. The compounds are understood to disrupt this protein–protein interaction by modifying the cysteine residues on Keap1, leading to the stabilization and activation of Nrf2 [15–16]. Interestingly, Hu et al identified (S, R, S)-1a (PDB IDs: 4L7B; 1VV) as the first cysteine-independent activator, functioning as a direct inhibitor of Keap1–Nrf2 binding [17–19]. A recent study has also explored the potential of phytochemicals acting as direct Nrf2 activators by targeting the Keap1–Nrf2 protein-protein interaction [20].

Gallic acid (3,4,5-trihydroxybenzoic acid) is a naturally-occurring polyphenol found in berries, tea, grapes, and medicinal plants. It is one of such compounds known for its potent antioxidant and cytoprotective effects [21]. Lee et al in their computational study suggests that gallic acid and other phytochemicals may interact with Keap1 and dissociate Nrf2 from it [20]. However, their findings have not been consistently validated with in vitro and in vivo antioxidant studies. In addition, there is a lack of studies that integrate computational predictions with biochemical evidence on the activities of antioxidant enzymes and levels of oxidative stress biomarkers in target tissues, after treatment with exogenous antioxidants such as gallic acid and ascorbic acid.

Therefore, this present study was designed to fill the gap by investigating the protective effects of gallic acid on enzymatic (SOD, CAT, GPx, GST) and non-enzymatic (GSH) antioxidant defenses, as well as on oxidative stress biomarkers (MDA, PCO, NO, PC) in the heart, erythrocytes, liver, kidney, femur, and spleen of mice subjected to benzene toxicity. We also evaluated its effects on hematological parameters in their whole blood. Beyond the direct scavenging of ROS, we further hypothesized that gallic acid may activate Nrf2 signaling by interfering with Keap1-Nrf2 complex formation. To test this hypothesis, molecular docking was utilized to assess the binding affinity and interactions of gallic acid within the Kelch-domain binding pocket of Keap1. This integrated computational, in vitro, and in vivo approach aimed to provide additional insights into the mechanism and ability of gallic acid in protecting the antioxidant defense system against chronic oxidative stress, as well as its potential to inhibit Keap1–Nrf2 binding, prevent Nrf2 degradation, and enhance its nuclear translocation, in a process that ultimately promotes cellular resilience against benzene-induced toxicity.

## Methods

### Chemicals and reagents

Benzene, gallic acid, ascorbic acid, and sesame oil were products from Tokyo Chemical Industry Co. Ltd, Tokyo, Japan. Assay kits used for antioxidant enzymes and oxidative stress biomarkers assay were products of Randox Laboratory Ltd, Ardmore, Co., Antrim, UK. Other chemicals used were obtained commercially and of analytical grade.

### Experimental animals

Thirty-six (36) Swiss male mice, about 4 months old and with an average weight of 27 ± 2 g, were used for the experiment. The mice were obtained from the Animal Holding Unit of the Department of Biochemistry, University of Ilorin, Ilorin, Nigeria, and were acclimatized for 14 days under standard housing conditions (temperature, 25 ± 2 ⁰C; humidity, 50-70%; 12-hour light/dark cycle). They were given free access to clean water and a standard commercial diet (Topfeed, Premier Feed Mills Co. Ltd., Nigeria) *ad libitum*.

### Ethical clearance

Ethical clearance on the use of laboratory animals was issued by Ethics Committee of University of Ilorin, Ilorin, Nigeria and we adhered strictly to the Principles of Laboratory Animal Care (NIH Publication, No. 85-23) [22].

### Experimental study design

The mice were randomized into 6 groups of 6 animals each. The grouping was done as follows:

Group A (Normal Control): 0.2 ml/20 g distilled water + 0.2 ml/20 g sesame oil.

Group B (Negative Control): 150 mg/kg body weight benzene + 0.2 ml/20 g distilled water.

Group C: 150 mg/kg body weight benzene + 50 mg/kg body weight ascorbic acid (reference drug).

Group D: 150 mg/kg body weight benzene + 25 mg/kg body weight gallic acid. Group E: 150 mg/kg body weight benzene + 50 mg/kg body weight gallic acid. Group F: 150 mg/kg body weight benzene + 100 mg/kg body weight gallic acid.

Doses of benzene, sesame oil, gallic acid, and ascorbic acid were sourced from literature [23–25].

### Induction of benzene toxicity

Benzene toxicity was induced in mice via oral administration of benzene after it was diluted in sesame oil, in a single dose of 150 mg/kg body weight, once daily, 6 days/week, for 14 days. Ascorbic acid and gallic acid were also administered the same way simultaneously with benzene throughout the duration.

### Blood sample collection

At the end of 14 days (24 hours after the last treatment), the animals were anaesthetized with diethyl ether and venous blood via jugular vein was collected into EDTA bottles.

### Hematological test

To determine the level of toxicity, white blood cell count (WBC), red blood cell count (RBC), hemoglobin concentration (HGB), hematocrit (HCT), mean corpuscular volume (MCV), mean corpuscular hemoglobin (MCH), mean corpuscular hemoglobin concentration (MCHC), platelet count (PLT), and lymphocyte count (LYM) of the blood samples were analyzed immediately upon collection, by an automated hematological analyzer (SMT-50 Hematological Analyzer; Chengdu Seamaty Technology Co., LTD, Chengdu, China).

### Blood sample preparation

Whole blood sample was centrifuged at 2147 g for 10 minutes at 4 °C and plasma was carefully separated. The buffy coat, which contains leukocytes and platelets, was carefully aspirated with a pipette to minimize contamination. The remaining red blood cells (RBCs) were washed twice with 4 ml of phosphate buffer (100 mM, pH 7.4) to eliminate residual plasma proteins and other cellular components. After the second washing, supernatants were discarded to obtain packed RBCs. All samples were stored at -80 °C until needed for further analyses, following established RBC processing protocols [26–27].

### Red blood cell lysis

A total of 10 ml of 10x RBC lysis buffer was added to 1 ml of packed RBCs, and the mixture was incubated for 10-15 minutes at room temperature. The turbidity of the suspension was monitored until it disappeared, indicating lysis was successful. The lysate was then centrifuged at 1643 g for 5 minutes, after which the supernatant was pipetted out and frozen at - 80 °C until needed for further analyses [28–29].

### Preparation of tissue homogenate

The mice were quickly dissected and their organs (heart, liver, kidney, femur, and spleen) excised, cleansed of blood stains with cotton wool, weighed, and homogenized separately in ice-cold 0.25 M sucrose solution (1:5, w/v). The homogenates were centrifuged at 1643 g for 5 minutes to obtain the supernatants which were pipetted out and frozen at -80 °C until needed for further analyses [29].

## Biochemical analysis

### Determination of oxidative stress markers

#### Nitric oxide (NO)

Nitric oxide concentration in all tissue homogenates and erythrocyte lysates was estimated using the Griess reaction [30].

#### Principle

The assay is based on the reduction of nitrate to nitrite by nitrate reductase, followed by the reaction of nitrite with Griess reagent to produce a chromophore. Nitric oxide levels were expressed as µmol nitrite/mg protein.

#### Procedure

50 µl of each sample was incubated with 100 µl of Griess reagent at 25 ° C for 10 minutes. The absorbance was measured at 550 nm, and nitrite concentration was calculated using a sodium nitrite standard curve.

#### Malondialdehyde (MDA)

MDA concentration was determined using the method of Hunter (1963) as modified by Gutteridge and Wilkins (1982). Results were expressed as µmol MDA/mg protein [31].

#### Principle

MDA reacts with 2-thiobarbituric acid (TBA) under acidic and high temperature conditions to form a pink-colored product.

#### Procedure

0.5 ml aliquot of sample was mixed with 2.5 ml of TBA reagent (1 g of TBA dissolved in 100 ml of 0.2% NaOH and mixed with 3 ml glacial acetic acid). The mixture was vortexed thoroughly and incubated in a boiling water bath for 15 minutes. After cooling, the samples were centrifuged at 1207 g for 10 minutes, and the absorbance was read at 532 nm.

##### Calculation

MDA concentration was determined using the molar extinction coefficient of the MDA-TBA complex: MDA (µmol/mg protein) = (A_532_ / 1.56 × 10^6^) / protein concentration (mg/ml).

### Protein Carbonyl (PCO)

PCO concentration in the samples was determined by a spectrophotometric method described by Graziano *et al.,* 2015 [32].

#### Principle

This assay is based on 2,4-dinitrophenylhydrazine (DNPH), also known as Brady’s reagent, which is a specific probe that is able to react with protein carbonyl groups to form stable protein conjugated with dinitrophenylhydrazones (DNP). PCO concentration was expressed as µmol/mg protein.

#### Procedure

100 µl aliquot of the sample was mixed with 100 µl of 10 mM DNPH in an Eppendorf tube and incubated in the dark for 1 hour. After incubation, 100 µl of Trichloroacetic acid (TCA) was added and the mixture was centrifuged at 8586 g for 3 minutes. The supernatant was discarded, and the pellet was washed three times with 500 µl ethanol: ethyl acetate (1:1 v/v) to remove free DNPH. The final pellet was dissolved in 500 µl of 6 M guanidine hydrochloride and incubated for 10 minutes at 37 °C, followed by centrifugation at 8586 g for 3 minutes. A 100 µl aliquot of the resulting supernatant was transferred to a cuvette, and absorbance was measured at 370 nm.

##### Calculation

PCO concentration was determined using the molar extinction coefficient: PCO (µmol/mg protein) = (A_370_ / 22000 × 10^6^) / protein concentration (mg/ml).

### Protein concentration (PC)

Protein concentration of all samples was determined using the Bradford method (Bradford, 1976) and expressed as mg/dl protein [33].

#### Principle

The assay is based on the binding of proteins to Coomassie blue dye under acidic conditions, which changes the dye from brown to blue. The color intensity is proportional to protein concentration.

#### Procedure

Samples containing proteins were obtained from the supernatants of tissue homogenates and erythrocyte lysates after centrifugation, as prepared for oxidative stress biomarker assays.

For the assay, 30 μl of each protein sample (in triplicate) was mixed to 30 μl of protein preparation buffer (0.01 M, pH 7.4), followed by the addition of 1.5 ml of Bradford reagent. The mixture was incubated at room temperature for 5 minutes and absorbance was measured at 595 nm.

A standard curve was prepared using serial dilutions of bovine serum albumin (BSA) as the protein standard (0, 2.5, 5, 7.5, 10, 12.5, 15, and 20 µg in final volumes of 20 µl deionized water). 2, 5, and 10 µl sample protein solutions were diluted in 1 M NaOH to a final volume of 10 µl. Bradford reagent (1 ml) was added to both standards and samples, vortexed, and incubated for 5 minutes at room temperature. Absorbance was measured at 595 nm.

## Determination of non-enzymatic antioxidant

### Reduced glutathione (GSH)

The concentration of reduced glutathione in tissue homogenates and erythrocyte lysates was estimated according to the method of Sedlak and Lindsay (1968) [34].

#### Principle

The assay is based on the reaction of reduced glutathione (GSH) with 5,5’-dithiobis-(2-nitrobenzoic acid) (DTNB, Ellman’s reagent) which produces a yellow-colored 2-nitro-5-thiobenzoic acid (TNB). The total GSH content was expressed as µmol/mg protein.

#### Procedure

To 1 ml of the sample suspension (1 mg protein/ml), 1 ml of 10% Trichloroacetate (TCA) containing 1 mM EDTA was added to precipitate proteins. The mixture was centrifuged at 839 g for 10 minutes. 1 ml of the clear supernatant was then mixed with 0.5 ml of Ellman’s reagent and 3 ml of phosphate buffer (0.2 M, pH 7.4). The absorbance of the solution was measured at 412 nm.

## Determination of enzymatic antioxidant

### Superoxide dismutase (SOD)

A method originally described by Misra and Fridovich, 1972 was employed [35].

#### Principle

The principle of this assay is based on the ability of SOD to inhibit the autoxidation of epinephrine to adrenochrome at alkaline p^H^. One unit of SOD activity was defined as the amount of enzyme that inhibits 50% of the rate of epinephrine autoxidation, and the activity was expressed as U/mg protein.

#### Procedure

To 2 ml sample, 2.5 ml of carbonate buffer (0.05 M, pH 10.2) was added and equilibrated at room temperature. The reaction was initiated by adding 0.3 ml of freshly prepared epinephrine solution (30 mM). The increase in absorbance was recorded at 480 nm for 4 minutes against a reference solution without the enzyme sample.

### Catalase (CAT)

Catalase activity was determined according to the procedure of Sinha [36].

#### Principle

In this assay, dichromate in acetic acid is reduced to chromic acetate when heated in the presence of H_2_O_2_ per minute, with perchromic acid formed as an unstable intermediate. The chromic acetate so formed is measured spectrophotometrically. One unit (U) of CAT activity was defined as the amount of enzyme required to decompose 1 µmol of H_2_O_2_ per minute under assay conditions. Results were expressed as U/mg protein.

#### Procedure

The assay mixture contained 0.5 ml of H_2_O_2_ (0.2 M), 1.0 ml of phosphate buffer (0.01 M, pH 7.4), and 0.4 ml of distilled water. The reaction was initiated by adding 0.2 ml of the samples as the enzyme source. At 0, 30, 60, and 90 seconds of incubation, 2 ml of dichromate-acetic acid reagent was added to stop the reaction. Control tubes received 2 ml of homogenate or lysate after the addition of the acid reagent. All tubes were then heated for 10 minutes to develop the green chromic acetate color, and absorbance was measured at 570 nm against distilled water.

### Glutathione Peroxidase (GPx)

GPx activity in each sample was determined in triplicate using the method of Paglia and Valentine (1967) [37].

#### Principle

The assay is based on the GPx-mediated oxidation of reduced glutathione (GSH) during the reduction of cumene hydroperoxide substrate. In the presence of glutathione reductase and NADPH, the oxidized glutathione (GSSG) is immediately converted back to GSH accompanied by the oxidation of NADPH to NADP^+^. The decrease in absorbance at 340 nm, corresponding to NADPH consumption is directly proportional to GPx activity. Since catalase can also degrade hydroperoxides and interfere with this assay, it is deactivated by the cyanide present in the Drabkin reagent. One unit (U) of GPx activity in each sample was defined as the amount of enzyme that catalyzes the oxidation of 1 nmol NADPH per minute. The activity of GPx is expressed as U/mg protein.

#### Procedure

Samples (5 ml) were diluted with 1 ml Ransel diluting agent and incubated for 5 minutes, followed by addition of 1 ml Drabkin reagent and mixing. For the assay, 0.1 ml of the diluted sample was combined with a solution containing 1 unit of glutathione reductase and 0.1 ml of 0.2 mM NADPH. Upon addition of 40 µl 1.5 mM cumene hydroperoxide solution, the absorbance was monitored at 340 nm over a 3 minutes period to determine the rate of conversion of the NADPH to NADP^+^.

### Glutathione S-tranferase (GST)

The determination of glutathione S-transferase activity was performed according to Habig *et al.,* (1974) using 1-chloro-2,4-dinitrobenzene (CDNB) as a substrate [38].

#### Principle

GST catalyzes the conjugation of L-glutathione (GSH) to CDNB. The reaction product, GS-DNB conjugate, absorbs light at 340 nm. The rate of increase in the absorption is directly proportional to GST activity in the sample.

#### Procedure

GST activity was assayed in the supernatant of tissue homogenates and erythrocyte lysates. Briefly, 180 µl GST buffer (9.8 ml of phosphate-buffered saline), 100 µl GSH, and 100 µl CDNB were added to 20 µl sample. The absorbance was measured at 340 nm for 5 minutes at 1-minute intervals. One unit (U) of GST activity was defined as the amount of enzyme that conjugates 1 µmol of CDNB with GSH per minute under assay conditions. Results were expressed as U/mg protein.

### Drug target preparation

The 3D crystal structure of human Keap1 (PDB ID: 4L7B) is a dimer (subunits A: PRO322-HIS612; B: ALA321-HIS612) that contains the Kelch domain of Keap1, in complex with compound (S,R,S)-1a inhibitor, a cyclohexane-carboxylic acid derivative (PDB ID: 1VV), retrieved in PDB format January 7, 2025, from Protein Data Bank [15,19]. The drug target was prepared by removing redundant subunit (A), acetate (ACT) and sodium (Na) ions, heteroatoms, and water molecules using PyMOL visualization tool [39]. The unique ligand compound (S, R, S)-1a, which served as one of the control ligands was extracted from the catalytic subunit B, in addition to its 3 other analogues (PDB IDs: 1VW, 1VX, 2FS) extracted from 4L7B-related structures (PDB IDs: 4L7C, 4L7D, 4N1B) respectively [15,19]. Both target and ligands were saved in PDB and SDF formats, respectively. Using PyMOL allows us to visualize and predict the grid co-ordinates around the Kelch domain binding pocket, while Discovery Studio visualizer [40] helps in identifying and characterizing the residues at this binding site.

### Virtual screening

DataWarrior software [41] was used to determine the drug-likeness and ADMET (absorption, distribution, metabolism, excretion, and toxicity) properties of Keap1 inhibitors (1VV, 1VW, 1VX, 2FS) in accordance with Lipinski’s rule of five [42–43]. In addition, the phytochemicals of interest, gallic acid (Pubchem ID: 370) and ascorbic acid (Pubchem ID: 54670067), were downloaded from the Pubchem database [44]. Using the “calculate compound properties from chemical structure” feature in Datawarrior, the compounds were further screened for mutagenicity, carcinogenicity, reproductive effectiveness, ligand efficiency, drug-likeness, and irritancy (Table 1) [45–47].

**Table 1:**
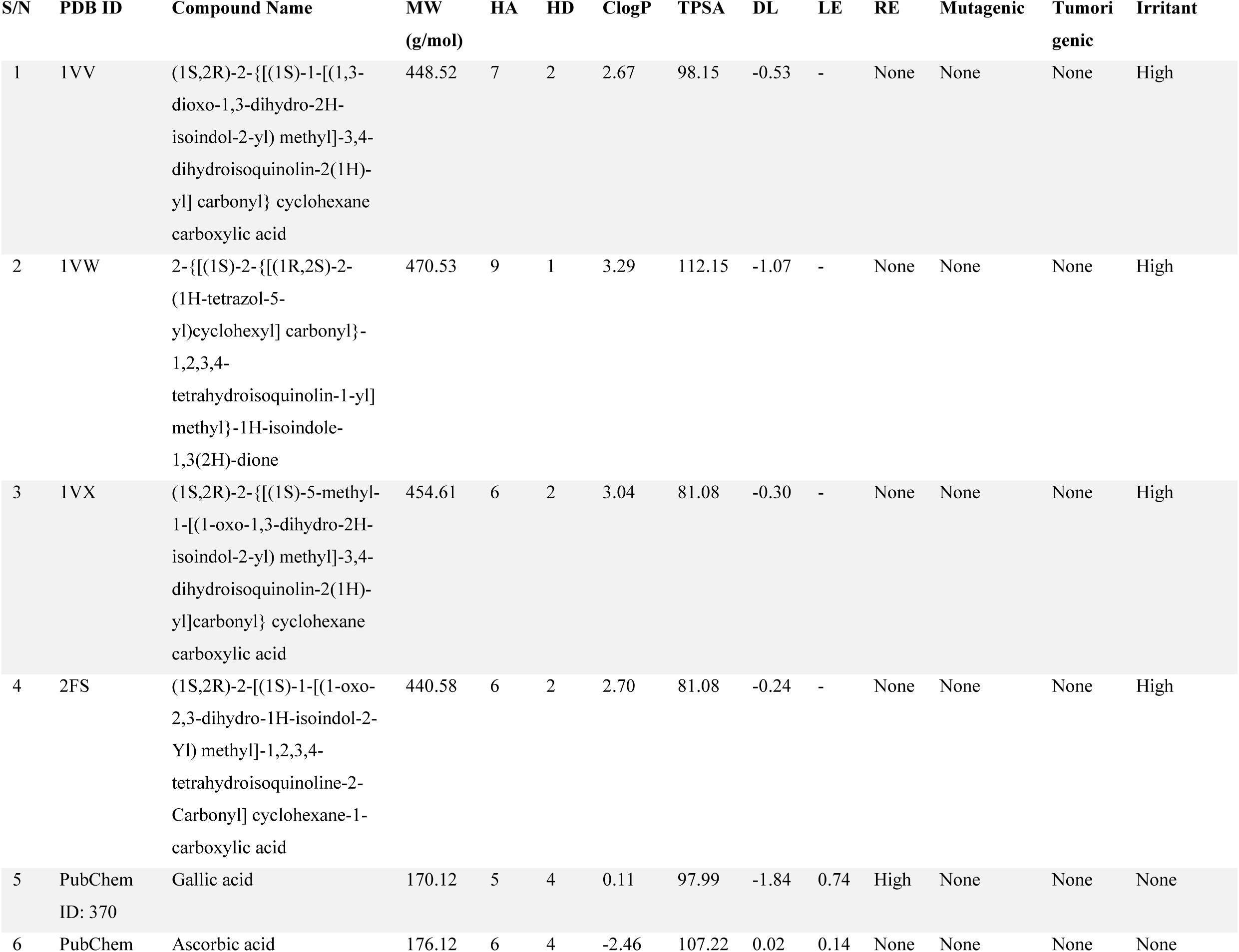
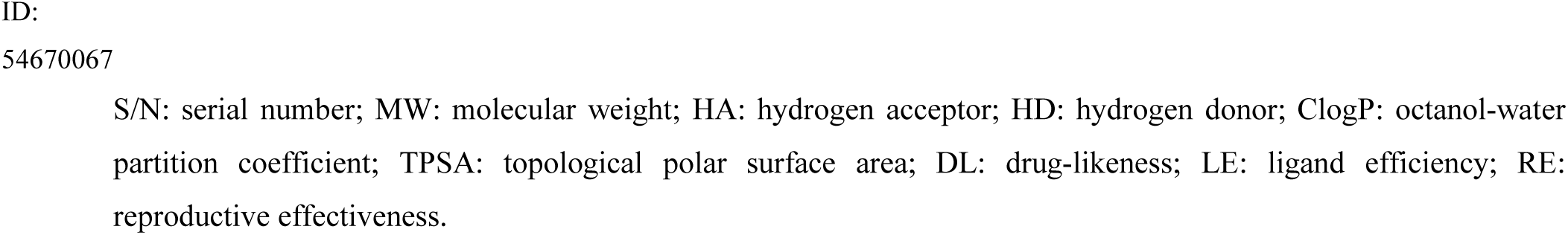
Drug-likeness and ADMET (Absorption, Distribution, Metabolism, Excretion, and Toxicity) properties of small molecule inhibitors of human Kelch-like ECH-associated protein 1 (Protein Data Bank ID: 4L7B)

### Molecular docking analysis

PyRx virtual screening tool [49] was used for the molecular docking. The prepared drug target Keap1 was loaded on PyRx in PDB format, hydrogen atoms were added to ensure the protein is correctly protonated and made as macromolecule, after which the screened compounds were imported in SDF format. These compounds were subjected to energy minimization using the optimization algorithm tool of PyRx, and the required force field was set at “ghemical”, adjusting the positions of atoms in the compounds in order to reduce their overall energy and steric clashes, to attain stable conformations. The compounds were converted to PDBQT format for compatibility with the docking algorithm Autodock Vina. Docking was performed specifically at the Kelch domain-binding pocket of the protein. The 3D docking grid box which encloses this region, where the compounds will bind was centered at co-ordinates (X: -28.0675, Y: 50.4914, Z: -37.1297) with grid box dimensions of 10.4482 × 12.2644 × 25.0877 Å, along the same axes, respectively. This type of docking is semirigid, where the structure of receptor (Keap1) remains rigid while the phytochemicals and control ligands have some degree of flexibility at the binding pocket. In the molecular docking, the PyRx AutoDock Vina Wizard exhaustive search docking function was used because of its balance between computational efficiency and accuracy [47–48]. To ensure the feasibility of the docking study protocol, 4 Keap1 analogue inhibitors (1VV, 1VW, 1VX, 2FS) were extracted from their respective Keap1 targets (PDB IDs: 4L7B, 4L7C, 4L7D, 4N1B). These inhibitors were redocked into the Kelch domain of 4L7B, before docking the phytochemicals at the same site. Their results were exported as PDBQT files and visualized using PyMOL and Discovery Studio to evaluate the best poses (binding modes), hydrogen bonding, hydrophobic interactions, and molecular fit within the binding pocket. Their binding energy scores were saved in excel format for statistical analysis. To generate the docking scores for the compounds, another round of docking was performed using the “Dock structure into protein cavity” feature on DataWarrior.

### Statistical analysis

All biological data were expressed as the mean of six determinations ± standard error of mean (SEM) or standard deviation (SD) for hematological parameters. Binding affinity scores were reported as the mean of nine determinations ± SEM. Statistical analysis of all data was performed using one-way analysis of variance (ANOVA), followed by Tukey’s multiple comparisons test as a posthoc test using Graphpad Prism version 8.0 (Graphpad Software Inc., California, USA). All data were considered statistically different at *P<.05*.

## Results

### Hematological parameters

Benzene exposure in untreated mice (Group B) caused a significant reduction in several hematological parameters such as WBC, RBC, HGB, HCT, PLT, and LYM. However, administration of gallic acid across the treatment groups (Groups D-F) reversed these observed alterations when compared with the normal control (Group A). The therapeutic effects of gallic acid at 50 and 100 mg/kg body weight (Groups E and F) were comparable to those of the standard antioxidant ascorbic acid (Group C). No appreciable changes were observed in the levels of MCV, MCH, and MCHC across all groups (Table 2).

**Table 2:**
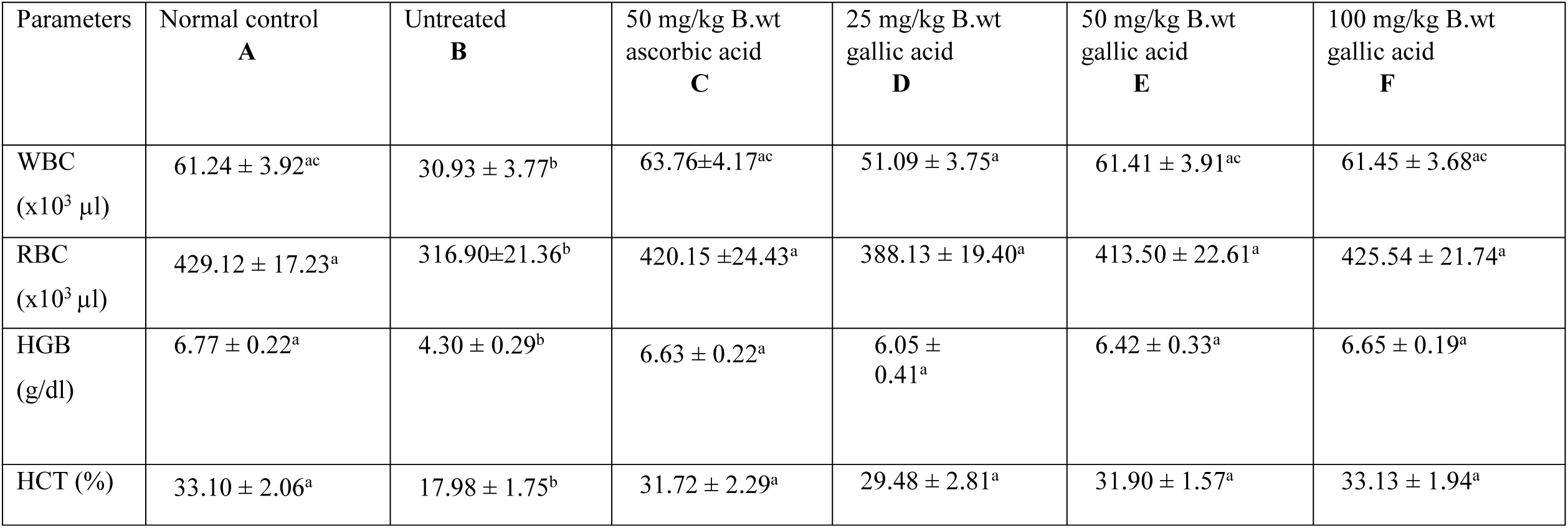
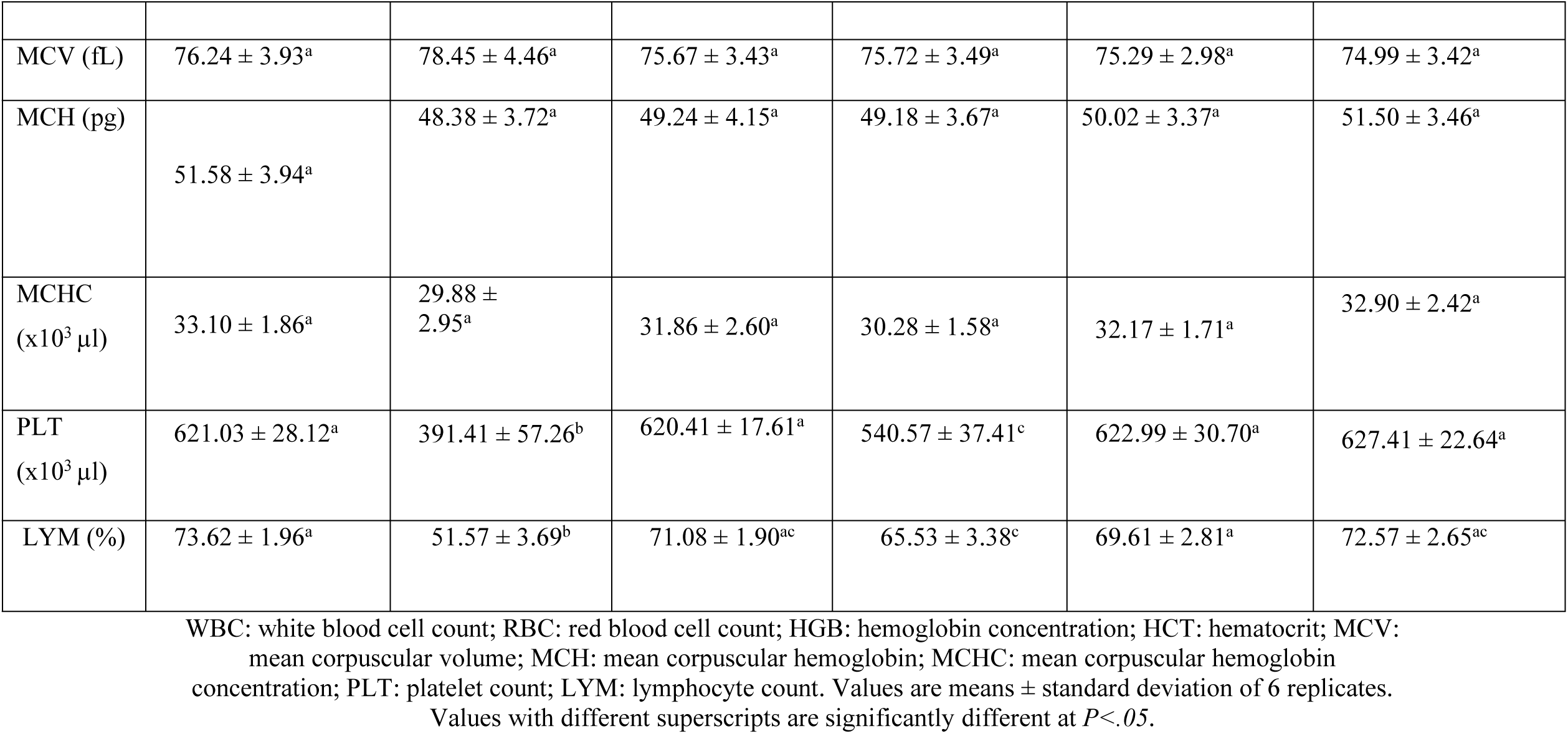
Effects of gallic acid on selected hematological parameters of mice with benzene-induced toxicity.

### Antioxidant enzymes and oxidative stress biomarkers

Biochemical assays in tissues of interest (heart, erythrocytes, femur, spleen, kidney, and liver) revealed that benzene exposure in untreated mice (Group B) significantly reduced the activities of selected antioxidant enzymes, including SOD, CAT, GPx, and GST. Treatment with gallic acid at different doses (Groups D-F) restored the activities of these enzymes to levels comparable with the normal control (Group A), similar to the effects observed with ascorbic acid (Group C) (Figures 1A-6A).

**Figure 1.**
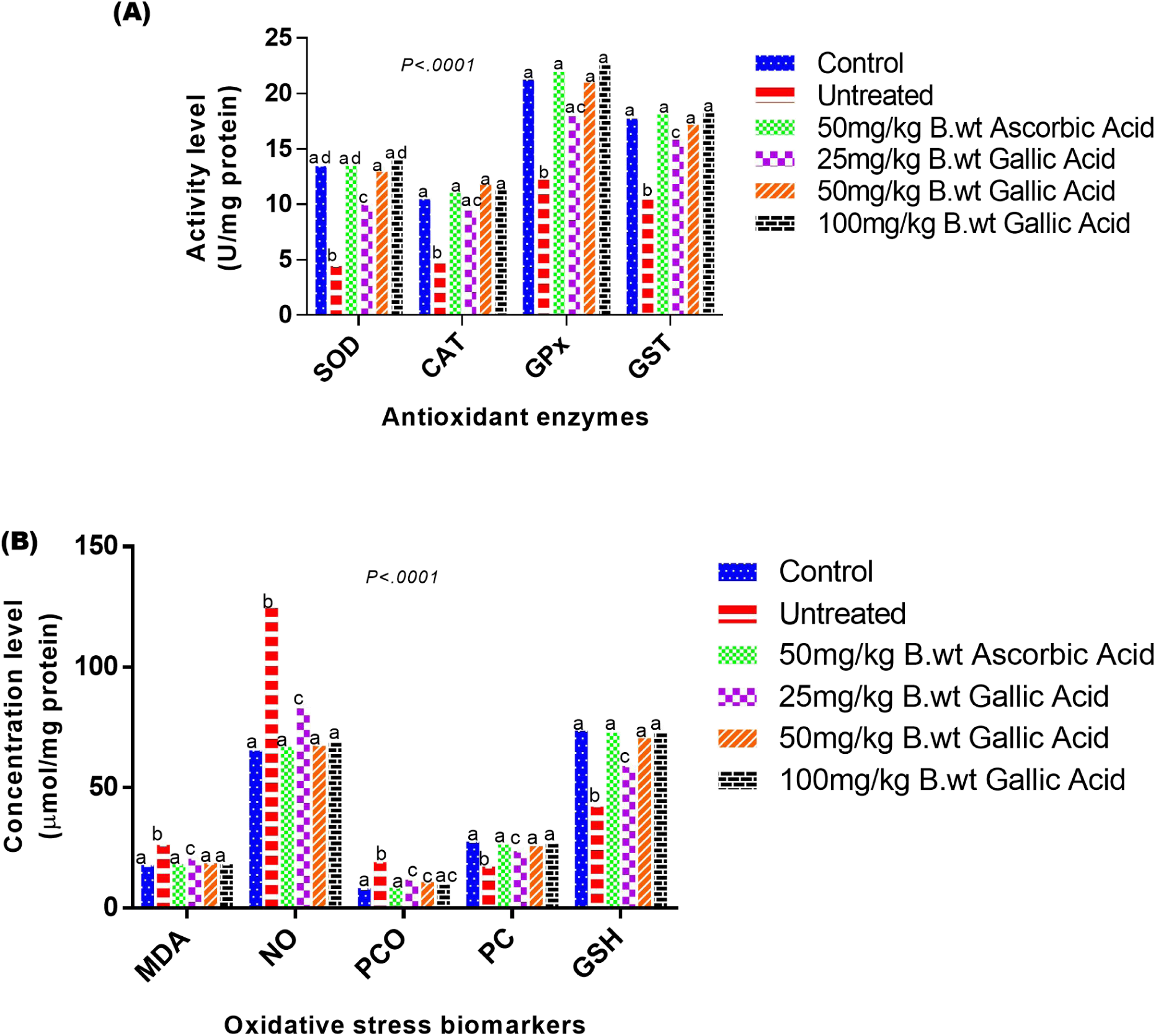
Effects of gallic acid on antioxidant enzymes (A) and oxidative stress biomarkers (B) in erythrocyte of mice with benzene-induced toxicity. Values are means ± SEM of 6 replicates. Bars with different superscripts are significantly different at *P<.05*.

**Figure 2.**
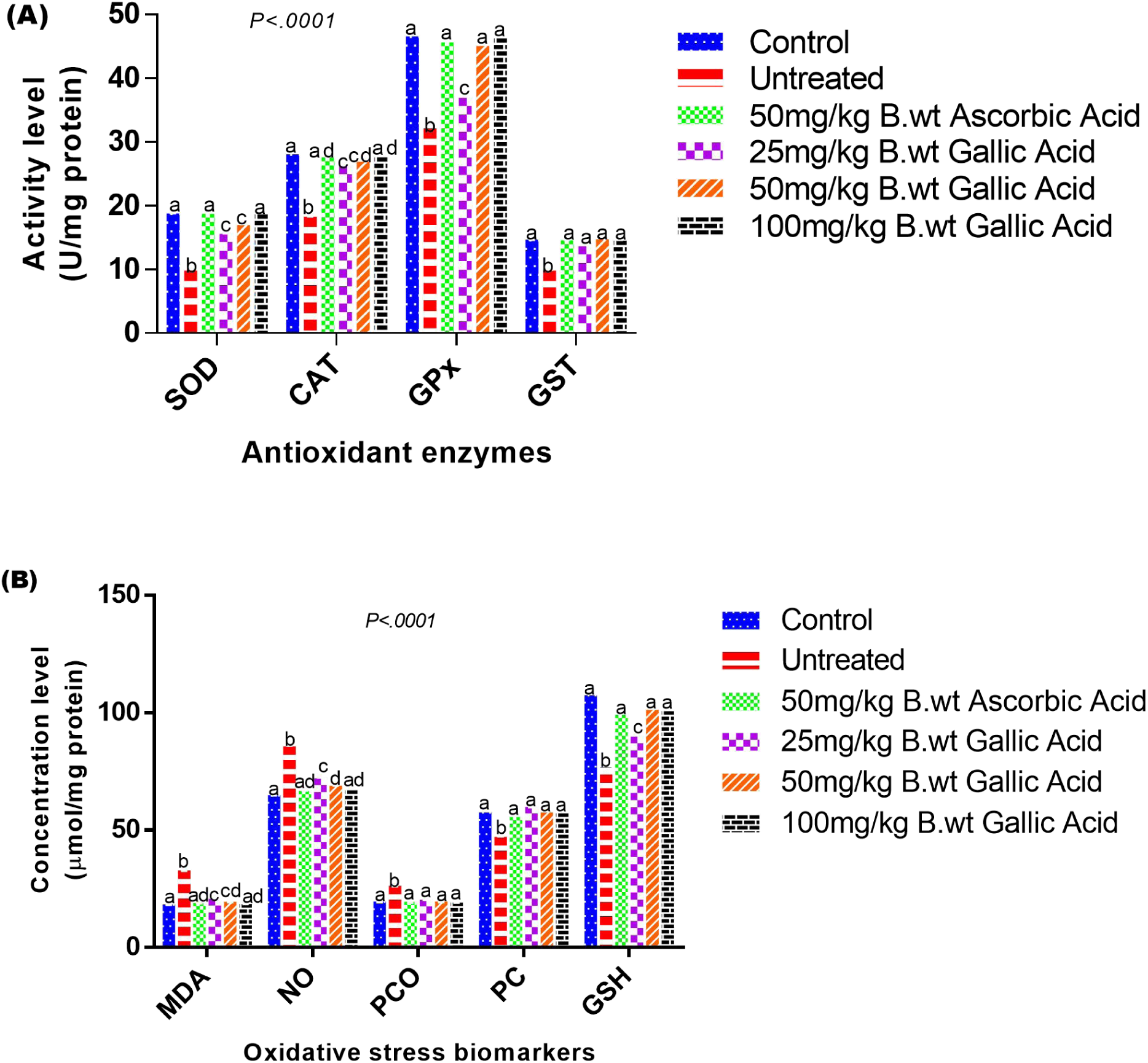
Effects of gallic acid on antioxidant enzymes (A) and oxidative stress biomarkers (B) in heart of mice with benzene-induced toxicity. Values are means ± SEM of 6 replicates. Bars with different superscripts are significantly different at *P<.05*.

**Figure 3.**
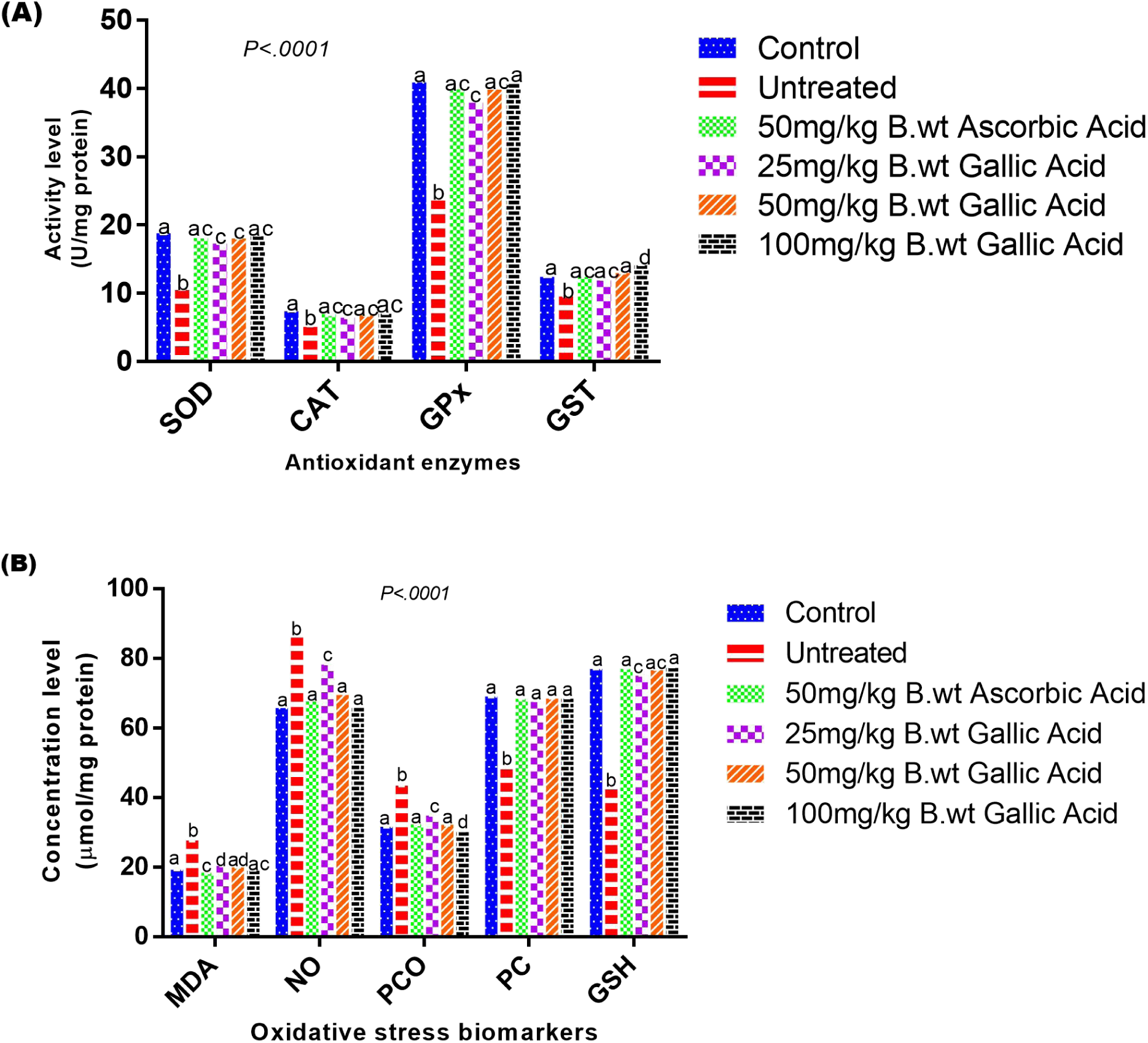
Effects of gallic acid on antioxidant enzymes (A) and oxidative stress biomarkers (B) in femur of mice with benzene-induced toxicity. Values are means ± SEM of 6 replicates. Bars with different superscripts are significantly different at *P<.05*.

**Figure 4.**
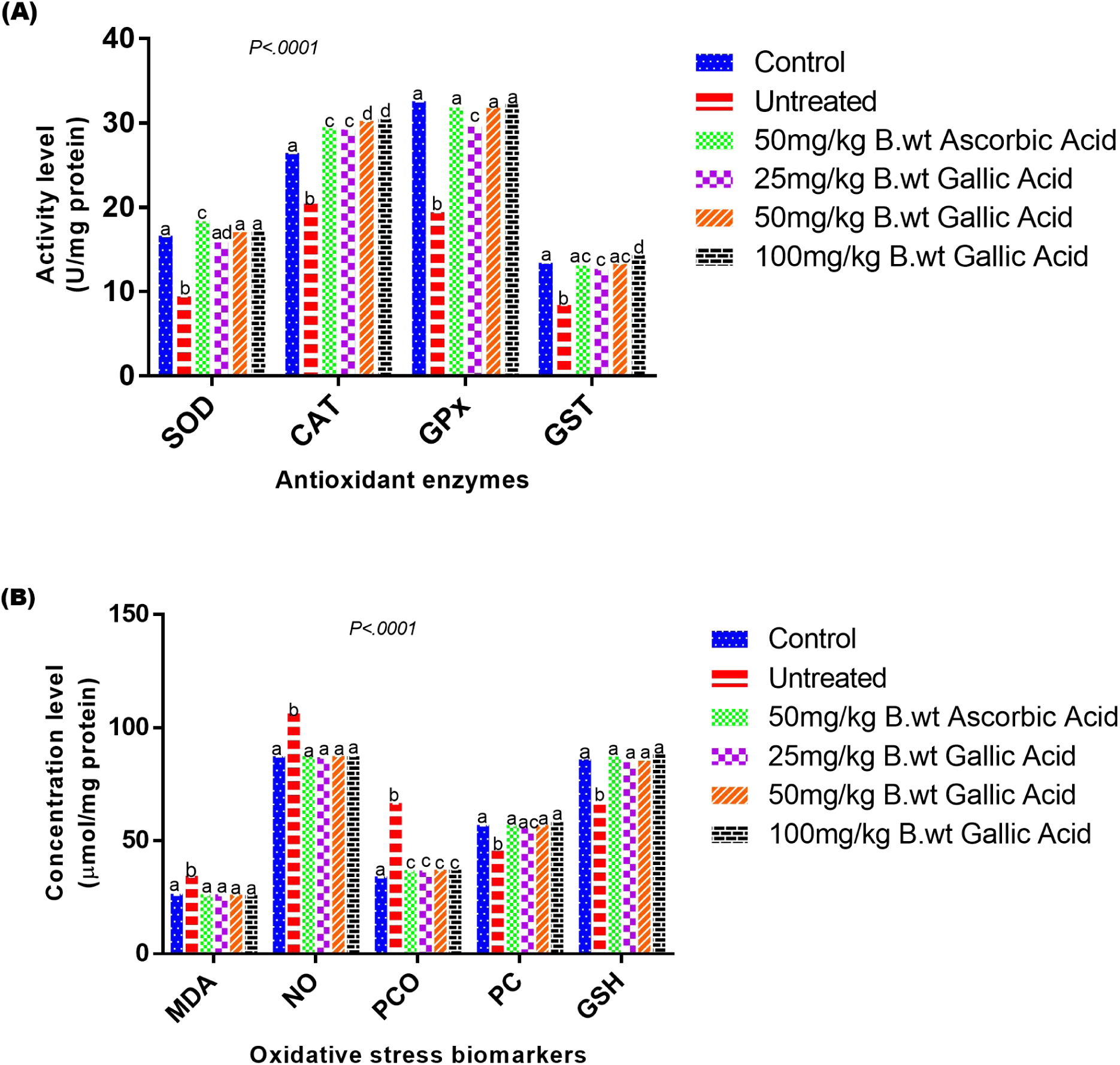
Effects of gallic acid on antioxidant enzymes (A) and oxidative stress biomarkers (B) in spleen of mice with benzene-induced toxicity. Values are means ± SEM of 6 replicates. Bars with different superscripts are significantly different at *P<.05*.

**Figure 5.**
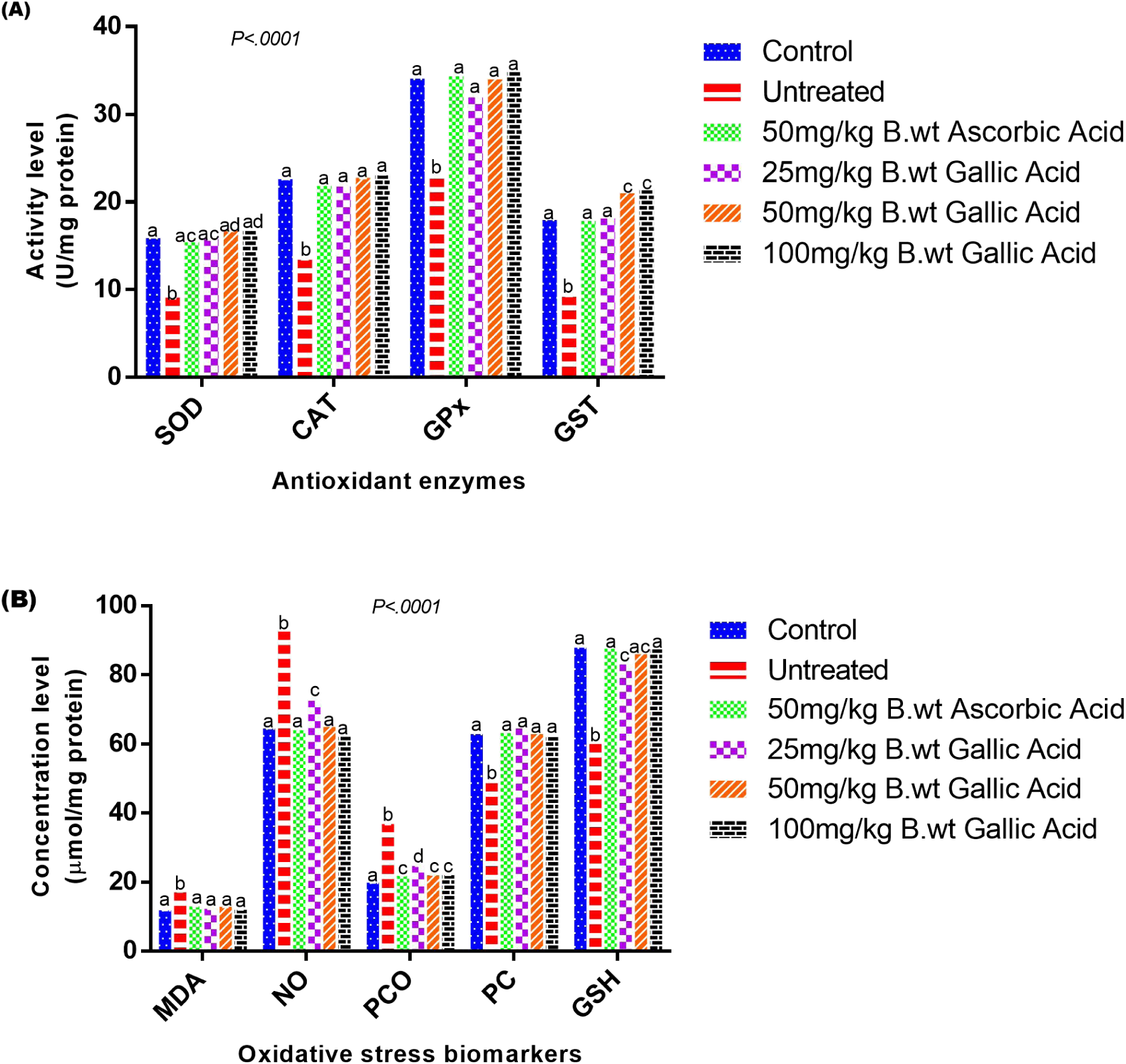
Effects of gallic acid on antioxidant enzymes (A) and oxidative stress biomarkers (B) in liver of mice with benzene-induced toxicity. Values are means ± SEM of 6 replicates. Bars with different superscripts are significantly different at *P<.05*.

**Figure 6.**
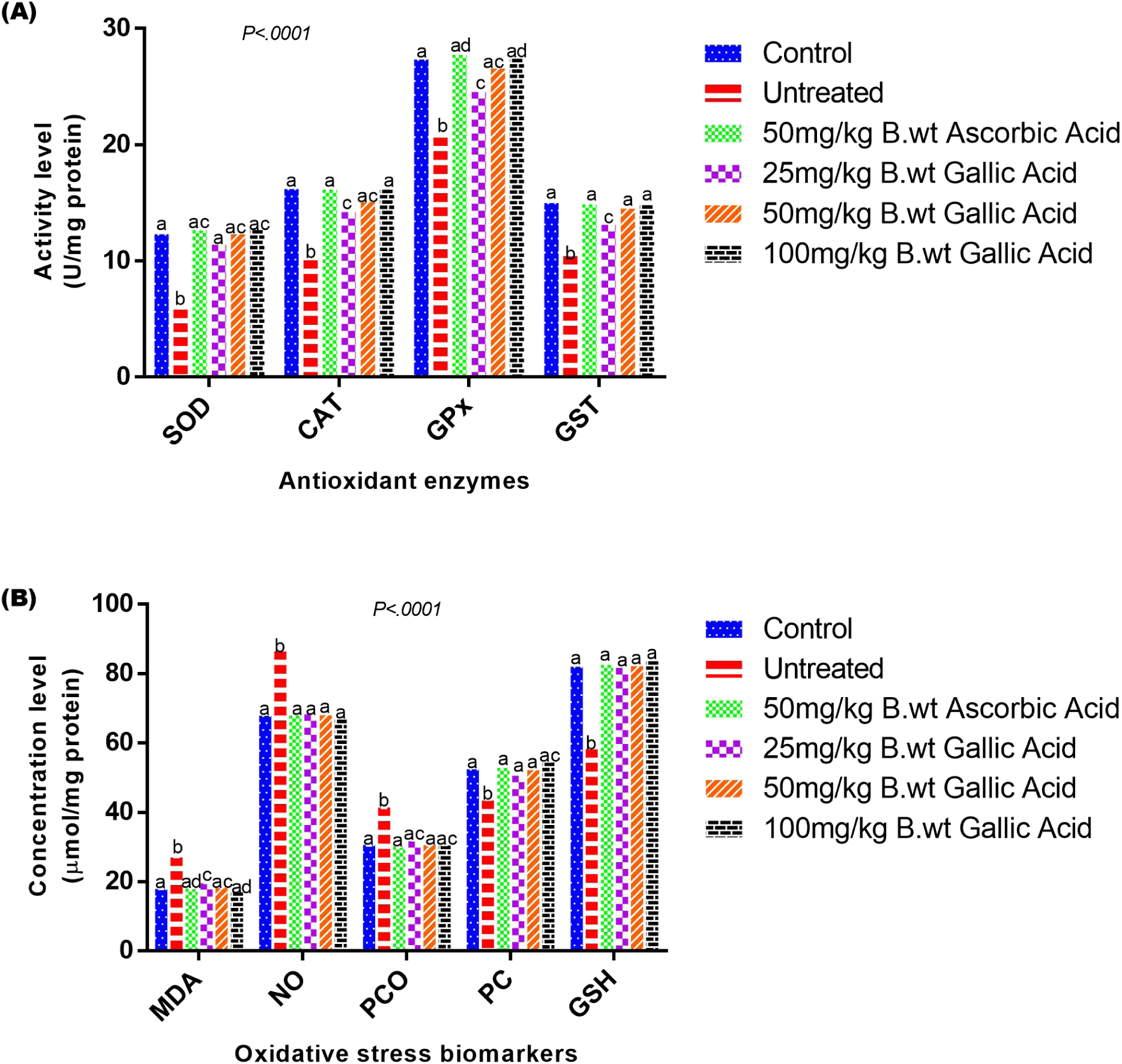
Effects of gallic acid on antioxidant enzymes (A) and oxidative stress biomarkers (B) in kidney of mice with benzene-induced toxicity. Values are means ± SEM of 6 replicates. Bars with different superscripts are significantly different at *P<.05*.

In contrast, untreated mice showed a significant increase in levels of oxidative stress biomarkers, such as MDA, PCO, and NO, along with significant reduction in the concentrations of GSH and PC across all tissues. However, administration of gallic acid at all tested doses reversed these alterations to near-normal levels, with results comparable to the ascorbic acid group (Figures 1B-6B).

### Molecular docking

To explore the potential mechanism of Keap1 inhibition by gallic acid, cysteine-independent inhibitors of Keap1 (control ligands; PDB IDs: 1VV, 1VW, 1VX, 2FS), gallic acid, and ascorbic acid were docked into the Kelch domain of the drug target. Docking results revealed that all control ligands, as well as gallic acid and ascorbic acid, exhibited high binding affinities and promising biochemical interactions with the Kelch domain of Keap1 (Table 3; Figures 7-8).

**Figure 7.**
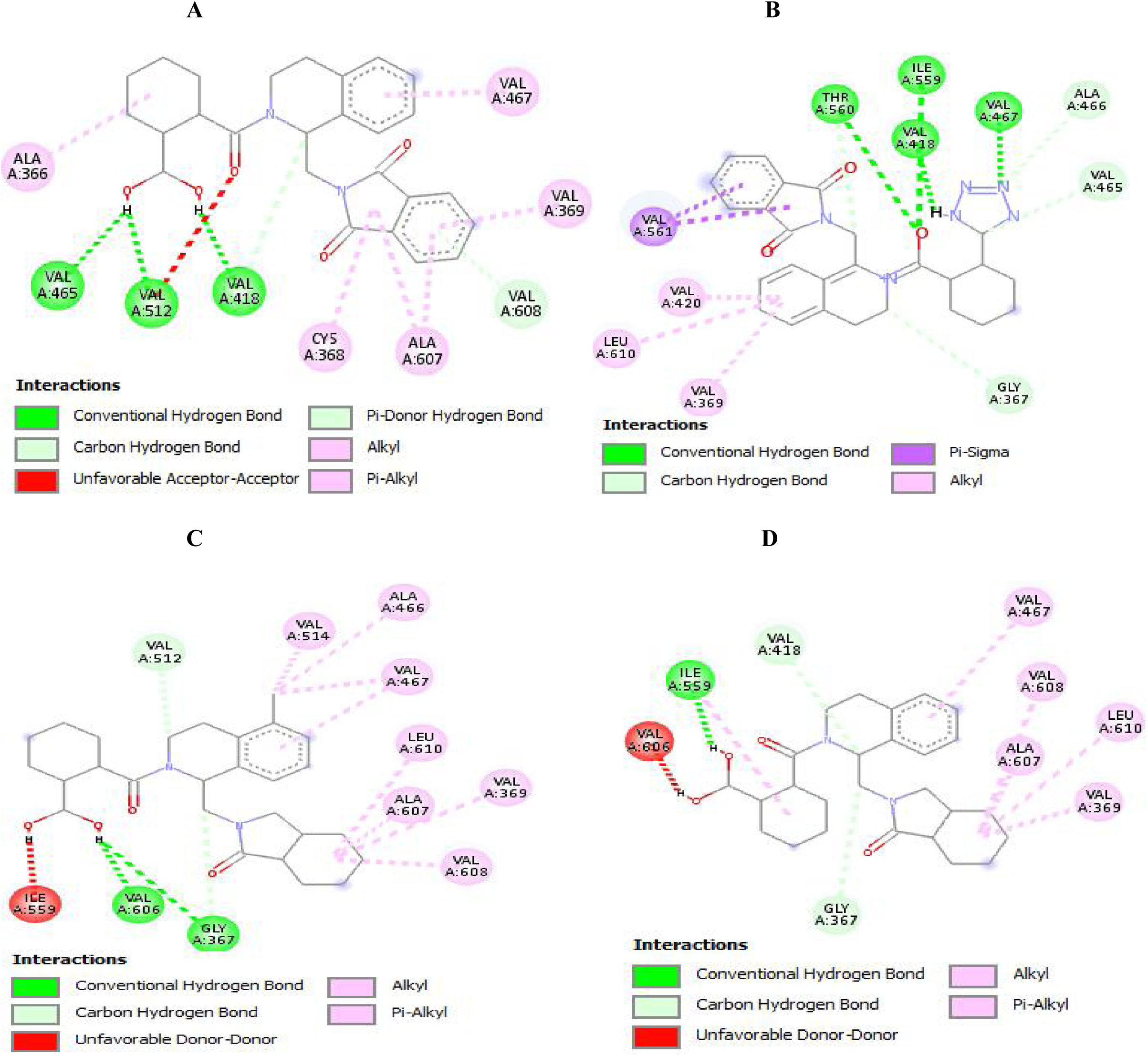
2D representation showing the binding interactions of (A) PDB ID_1VV pose 1, (B) PDB ID_1VW pose 1, (C) PDB ID_1VX pose 1, and (D) PDB ID_2FS pose 1 with Kelch domain binding pocket residues of human Kelch-like ECH-associated protein 1.

**Figure 8.**
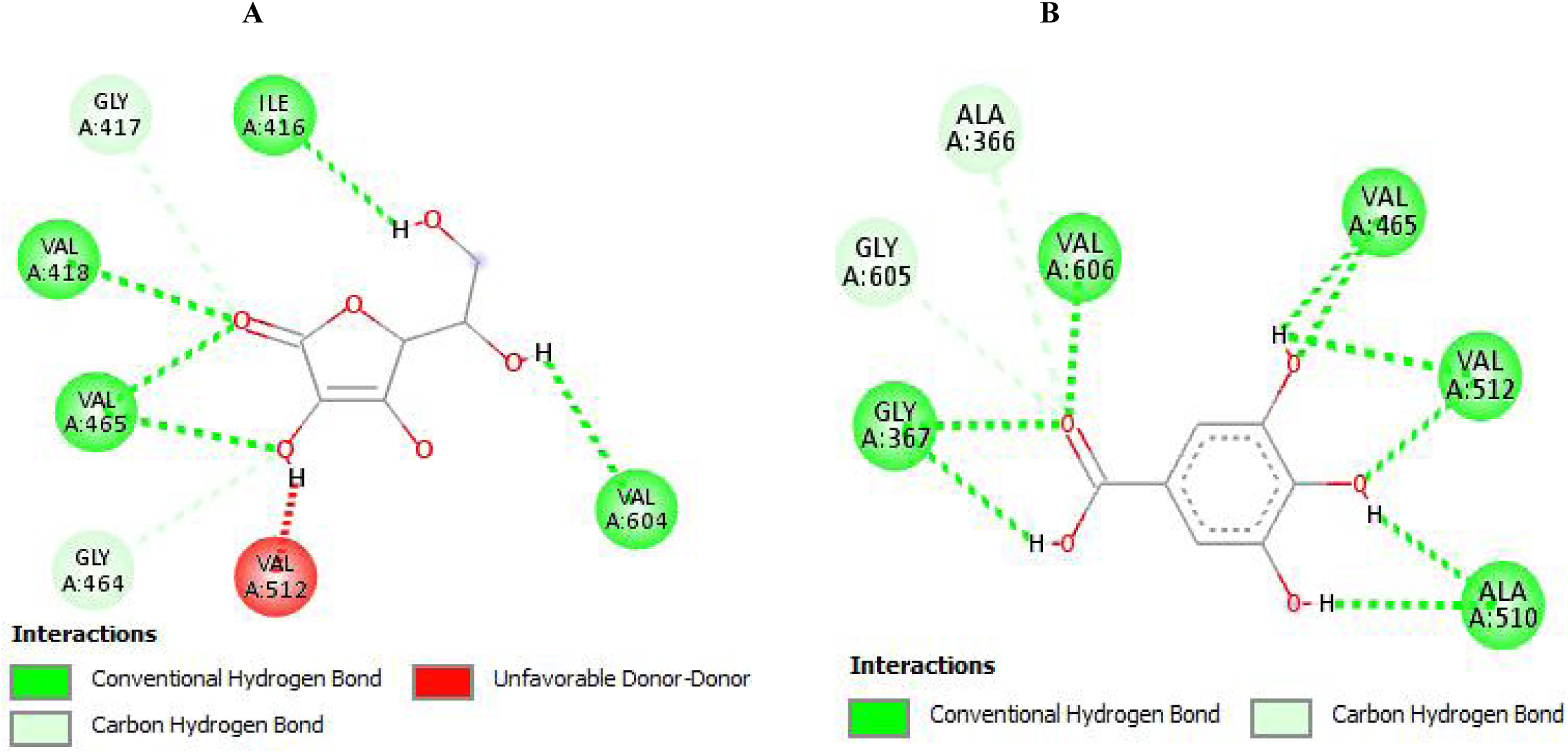
2D representation showing the binding interactions of (A) Ascorbic acid pose 2 and (B) Gallic acid pose 2 with Kelch domain binding pocket residues of human Kelch-like ECH-associated protein 1.

**Table 3:**
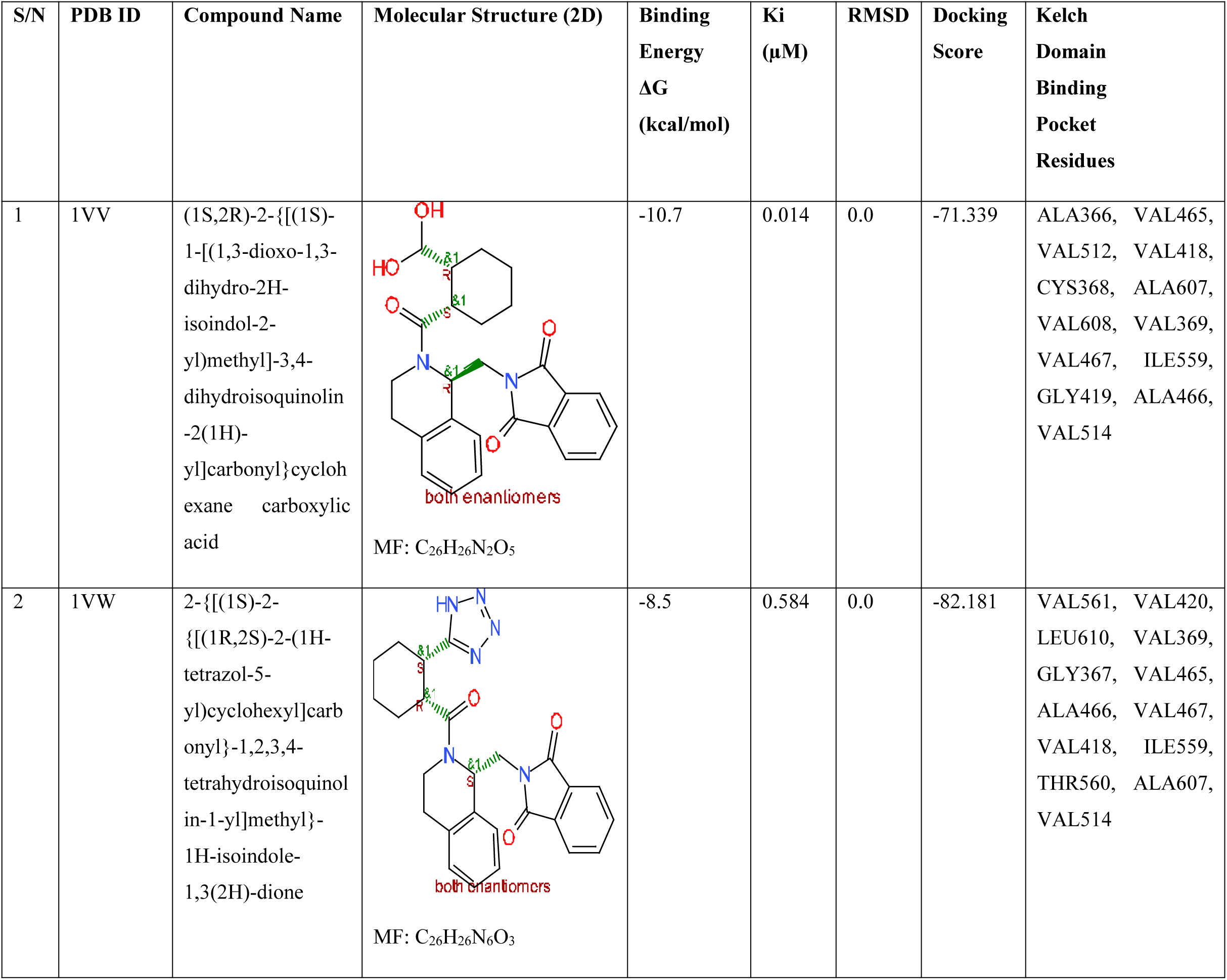
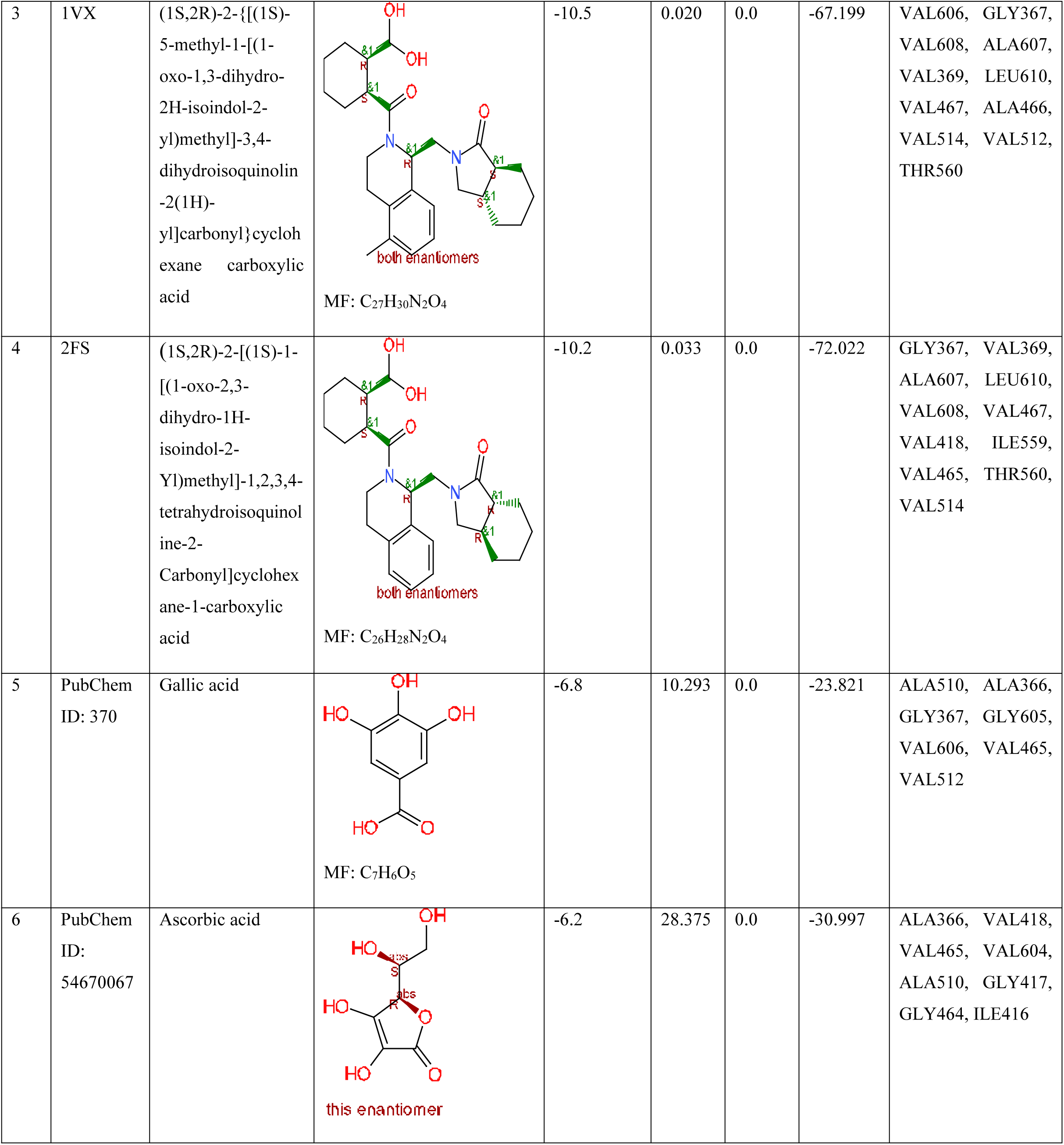
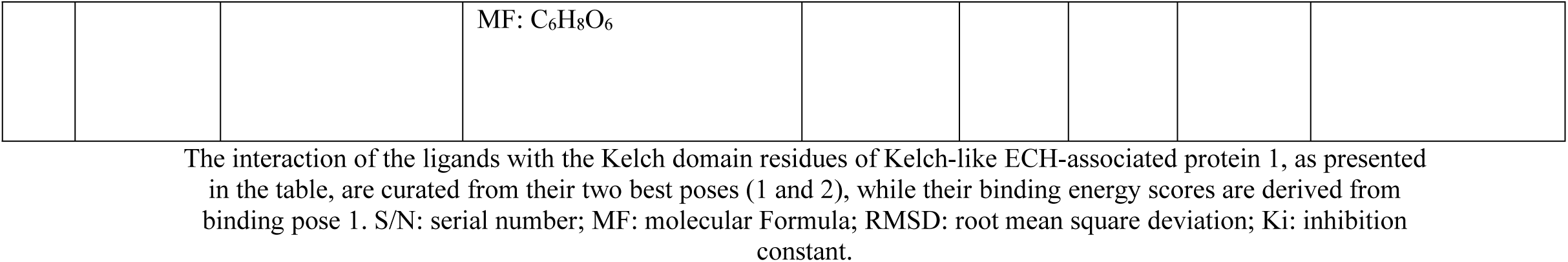
Molecular docking results of ligands binding at Kelch domain binding pocket of human Kelch-like ECH-associated protein 1 (Protein Data Bank ID: 4L7B)

In particular, gallic acid demonstrated a higher binding affinity (binding energy: -6.8 kcal/mol; 18 interactions) than ascorbic acid (-6.2 kcal/mol; 14 interactions) (Tables S5-S6 in Multimedia Appendix 1). However, both gallic acid and ascorbic acid showed comparatively weaker binding energies and docking scores relative to the higher molecular weight control ligands (Table 3; Tables S1-S4 in Multimedia Appendix 1).

## Discussion

### Effects of gallic acid on hematological parameters

In the untreated group, a significant decrease was observed in several hematological parameters, such as WBC, RBC, HGB, HCT, PLT, and LYM, whereas MCV, MCH, and MCHC showed no appreciable changes across all groups (Table 2).

Benzene-induced toxicity in humans usually occurs from occupational exposure. Benzene is metabolized primarily in the liver by cytochrome P_450_ 2E1 (CYP2E1), an isozyme of the cytochrome P_450_ mixed-function oxidases [7,67–68]. Benzene oxide, the first intermediate, is converted into phenol, hydroquinone, benzoquinone, catechol, and muconic acid or muconaldehyde. In the bone marrow, myeloperoxidase further oxidizes these phenolic metabolites to form free radicals capable of damaging hematopoietic tissues [2,69–70].

The harmful effects of these reactive metabolites in the bone marrow leads to multiple complications. For example, free radical-induced damage to myeloid progenitor cells can impair erythropoiesis and reduce lifespan of erythrocytes, resulting in varying degrees of anemia [6]. Furthermore, chemical-induced hemolytic anemia may occur when benzene metabolites disrupt cytoskeletal membrane proteins, leading to premature destruction of red blood cells [6]. Benzene toxicity has also been strongly associated with acute myeloid leukemia [71], supporting earlier studies by Aoyama who reported that mice exposed to 200 ppm benzene for 6 hours per day for 7 days, or 50 ppm for 14 days, exhibited decreased white blood cell counts in both blood and spleen [72]. This finding is consistent with our own observations in alterations of hematological parameters in benzene-exposed mice (Table 2). However, administration of gallic acid and ascorbic acid in the treatment groups reversed these alterations, restoring hematological parameters to levels comparable with those of the normal control.

### Effects of gallic acid on antioxidant enzymes

Both in vitro and in vivo analyses of this study revealed that benzene caused chronic oxidative stress in all examined tissues of untreated mice, as evidenced by significant reductions in the activities of antioxidant enzymes such as catalase (CAT), glutathione peroxidase (GPx), glutathione-S-transferase (GST), and superoxide dismutase (SOD) (Figures 1A-6A). This suggests that benzene, through its electrophilic metabolites, generated excessive free radicals that overwhelmed the enzymatic antioxidant defenses, the cell’s first line of protection against oxidative damage.

Deficiency in the activity of extracellular SOD and the resulting accumulation of superoxide radicals (O_2_^•−^), has been linked with hypertension due to higher conversion of nitric oxide (a vasodilator) to peroxynitrite, thus serving as a risk factor for cardiovascular disease [60]. Previous studies have also shown that during chronic oxidative stress, oxidants such as O_2_^•−^, H_2_O_2_, and ONOO^-^ can disrupt iron-sulfur (Fe-S) clusters of metalloproteins, especially those involved in antioxidant defense and redox reactions of electron transport in mitochondria, thereby inactivating them. The released iron can then react with H_2_O_2_ (formed via one-electron reduction of O_2_^•−^) in the so-called Fenton reaction, generating hydroxyl radical (^•^OH), an extremely reactive oxidant that indiscriminately damages whatever molecule it is next to in cells [8,61]. In addition, maintaining redox homeostasis is critical for cellular function; for example, studies have shown that type 2 diabetes mellitus may result from aberrant redox signaling mediated by these oxidants, leading to disruptions in redox balance [50].

However, our findings showed that administration of gallic acid significantly restored the activities of antioxidant enzymes, suggesting that gallic acid protects cells and their defense systems either by directly scavenging free radicals or by enhancing the performance of antioxidant enzymes to achieve optimal catalytic activity. At doses of 50 and 100 mg/kg body weight, gallic acid elicited the strongest protective effects, which compared favourably with the reference drug ascorbic acid, indicating comparable potency in mitigating benzene-induced oxidative stress. These observations align with earlier studies reporting that gallic acid can protect the cardiovascular system by inducing and sustaining antioxidant enzyme activity, thereby preventing the formation of advanced glycation end products [21,51–52].

### Effects of gallic acid on oxidative stress parameters

Analysis of oxidative stress biomarkers in the target tissues of untreated mice showed significant elevations in concentrations of malondialdehyde (MDA), nitric oxide (NO), and protein carbonyls (PCO), along with significant reductions in the level of reduced glutathione (GSH) and protein concentration (PC), compared to normal controls (Figures 1B-6B). These findings suggest that benzene exposure induced lipid peroxidation and protein oxidation. Free radical-mediated damage to biomolecules, when accumulated under acute or chronic oxidative stress, is a well-established contributor to the pathogenesis of a wide spectrum of diseases [8,53]. Increased levels of NO, MDA, and PCO further indicate that both reactive oxygen species (ROS) and reactive nitrogen species (RNS) substantially contributed to the oxidative stress observed in the target tissues.

Studies have demonstrated that NO, in the presence of excess superoxide radicals (O_2_^•−^), can generate peroxynitrite (ONOO^-^). In its protonated form (ONOOH), peroxynitrite acts as a potent oxidant, producing nitrogen (iv) oxide (^•^NO_2_) and hydroxyl radical (^•^OH). These highly reactive intermediates rapidly oxidize macromolecules, including membrane lipids, structural proteins, enzymes, and nucleic acids [8,54–55]. Meanwhile, oxidative stress-induced formation of lipid hydroperoxides is known to drive the conversion of low-density lipoprotein (LDL) cholesterol into its atherogenic oxidized form (OxLDL). OxLDL plays a central role in initiating inflammatory response, recruiting leukocytes to lesion sites, and promoting atherosclerosis through activation of smooth muscle cells and reduced NO bioavailability [56–57].

Similarly, elevated MDA and PCO levels have been linked with aberrant hydrogen peroxide (H_2_O_2_)-mediated redox signaling, contributing to the accumulation of advanced glycation end products implicated in diabetic complications and hypertension (50,58-59,61-62]. The reduced protein concentration observed in the target tissues suggests that protein integrity and function were compromised by oxidation or carbonylation, a known risk factor in Alzheimer’s disease (23). Likewise, depletion of GSH, being the most abundant non-enzymatic antioxidant in cells, indicates severe impairment of the redox-buffering system [63].

Nevertheless, gallic acid administration significantly lowered NO, MDA, and PCO levels, restoring them to levels similar to those of normal control. Also, GSH and protein concentrations were increased in groups with gallic acid treatment. These results demonstrate that gallic acid exhibited strong antioxidant effects, both by scavenging reactive species and by preserving endogenous antioxidant capacity. Our findings corroborate previous reports highlighting the role of gallic acid in mitigating oxidative stress-related complications in cancer, diabetes, cardiovascular disorders, and neurodegenerative diseases, due to its ability to restore and regulate redox homeostasis [64–66].

### Effects of gallic acid on Keap1

Molecular docking analysis revealed that gallic acid exhibited a strong binding affinity for the Kelch domain of Keap1, forming several polar and nonpolar interactions with amino acid residues including ALA^366^, GLY^367^, ALA^510^, GLY^605^, VAL^465^, Val^512^, and VAL^606^ (Table S6). This pattern of binding compares favorably with previously reported cysteine-independent small molecule inhibitors of Keap1 (PDB IDs: 4L7B, 4L7C, 4L7D, 4N1B) (Tables S1-S4).

Keap1 downregulates the transcription factor nuclear factor erythroid 2-related factor 2 (Nrf2), which is the principal regulator of cellular defense against oxidative stress. By interacting with the Kelch domain, gallic acid may disrupt the Keap1-Nrf2 complex, thus preventing proteasomal degradation of Nrf2. This predicted stabilization may allow Nrf2 to translocate into the nucleus, where it induces the transcription of genes encoding enzymes involved in both antioxidant defense and xenobiotic detoxification.

The predicted interaction of gallic acid with Keap1 is consistent with earlier reports showing that dietary polyphenols, such as quercetin and curcumin, can activate Nrf2 signaling through direct or indirect modulation of keap1 [73–75]. Similar to these compounds, gallic acid may function as a cysteine-independent inhibitor of Keap1, which helps explain the mechanism behind the enhanced antioxidant enzyme activities and the improvements in hematological and oxidative stress parameters observed in the target tissues of mice treated with gallic acid compared to the negative control.

### Conclusion

The findings of this study demonstrate that gallic acid can mitigate oxidative damage, enhance the antioxidant defense system, and potentially activate Nrf2 signaling by disrupting the Keap1-Nrf2 complex in the target tissues of mice exposed to benzene toxicity. Nevertheless, for antioxidant therapies to be more effective, whether in toxicological contexts or in disease prevention and management, future research should focus not only on scavenging free radicals but also on preventing the generation of highly reactive species at their source.

## Supporting information

Multimedia Appendix

## Acknowledgments

This study would not have been possible without the grace of God. We are also grateful to Prof. J.O. Adebayo for his guidance and supervision throughout this research.

## Authors’ Contributions

Conceptualization, TIO, BAA, MOO; Investigation, TIO, BAA, MOO; Data curation, TIO, BAA, MOO; Methodology, TIO, BAA, MOO; Formal analysis, TIO, BAA, MOO; Writing - original draft, TIO; Writing - review & editing, TIO, BAA; Software validation, TIO.

## Data Availability Statement

All data generated or analyzed during this study are included in its supplementary information files in Multimedia Appendix 2.

## Abbreviations

ADMET: absorption, distribution, metabolism, excretion, and toxicity
ARE: antioxidant responsive element
CAT: catalase
GPx: glutathione peroxidase
GSH: reduced glutathione
GST: glutathione S-transferase
Keap1: Kelch-like ECH-associated protein 1
MDA: malondialdehyde
Neh2: Nrf2-ECH homology 2
NO: nitric oxide
Nrf2: nuclear factor erythroid 2-related factor 2
PC: protein concentration
PCO: protein carbonyls
RNS: reactive nitrogen species
ROS: reactive oxygen species
SOD: superoxide dismutase

