## Supplementary figures and images for "Effects of Gallic Acid on Antioxidant Defense System and Nrf2 Signaling in Mice with Benzene-Induced Toxicity: In Vivo, In Vitro, and Computational Study"

### PyRx Screenshot.png

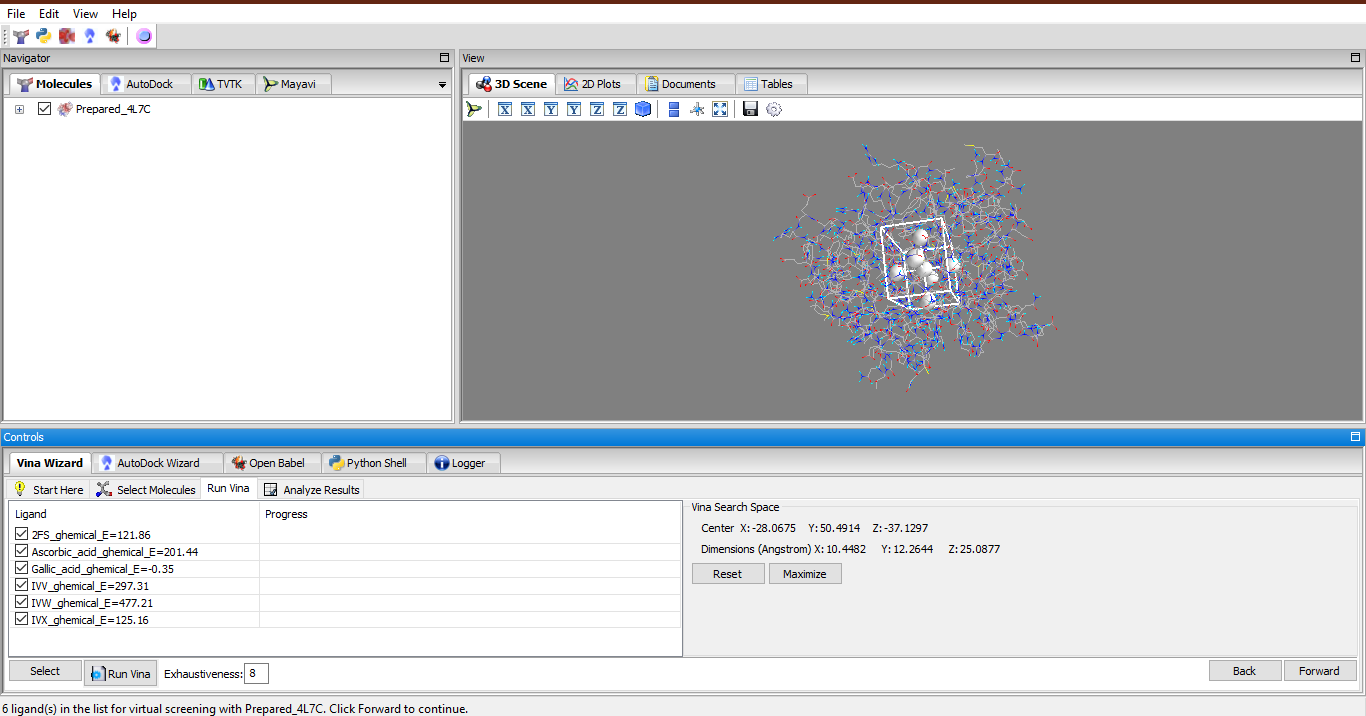
